# Cytosolic factors govern vimentin network architecture and mechanics

**DOI:** 10.64898/2026.01.08.697713

**Authors:** Dyuthi Sreekumar, Reza Shaebani, Mona Grünewald, Belinda König, A. S. Jijumon, Carsten Janke, Laura Schaedel

## Abstract

Vimentin intermediate filaments are key cytoskeletal components forming networks with architectures distinct from other intermediate filament types, enabling specialized functions. Although assembly of individual filaments from soluble subunits is well characterized, dissecting how vimentin networks are organized has been challenging, as existing in vitro systems do not mimic the structures observed in cells. Thus, how cells establish higher-order vimentin organization remains unclear.

Here, we reconstitute cell-like vimentin networks in vitro, using purified vimentin and extracts from mammalian and non-mammalian cells. Systematic variation of parameters reveals that cytosolic biomolecules, rather than intrinsic filament properties or generic ionic components, are the primary determinants of network architecture and mechanical behaviour. Importantly, network architecture is not universal but varies strongly according to the cell type from which the extract is derived, indicating that vimentin assemblies are tailored in a cell type-specific manner.

Thus, this extract-based reconstitution system enables mechanistic dissection of intermediate filament regulation under near-native biochemical conditions, bridging the current gap between purified systems and the cellular environment. Our findings show that vimentin architecture and mechanics emerge primarily from cytosolic biomolecular factors that organize filaments into cell type-specific networks. These results establish cytosolic regulation as a central mechanism specifying intermediate filament network architecture and function.

## Introduction

Intermediate filaments are key cytoskeletal elements implicated in a wide range of cellular processes and human diseases [1], [2], [3]. Among them, vimentin – the most abundant intermediate filament type – is important for directed cell migration [4], [5], [6], [7], force transmission and protection against large deformations [8], [9], [10], [11], [12], structuring the intracellular space and positioning of organelles [13], [14], and the regulation of cell identity through its roles in cell differentiation and tissue homeostasis [15].

While the exceptional mechanical properties and assembly dynamics of single vimentin filaments are well characterized [16], [17], [18], [19], major gaps remain in our understanding of how vimentin achieves its diverse roles, and how this differs from other intermediate filament types. In particular, a persisting challenge is to explain how vimentin filaments are organized into complex, cell-type-specific network architectures that strongly differ from other intermediate filaments [20], [21], [22], [23] – such as keratins, which typically form dense or strongly bundled meshworks, or desmin, which is organized along the sarcomeric structure of muscle cells – and how vimentin organization at the network scale governs its mechanical behavior and cellular functions.

This represents a fundamental knowledge gap compared to actin and microtubule cytoskeletal networks, where the relationship between architecture and function is well established [24], [25], [26], [27]. Progress in addressing these questions has been limited by the challenges of studying vimentin in the cellular context, where the dense and heterogeneous cytoplasm complicates the disentanglement of individual contributions of interacting cytoskeletal systems[28], [29], [30]. In vitro approaches have provided essential insights into the properties of vimentin assemblies based on non-specific interactions between filaments induced mainly by mono- and divalent ions and filament entanglement [31], [32], [33], [34], [35]; yet, these assemblies do not reproduce the well-organized, hierarchically structured networks found in cells. As a consequence, the critical link between vimentin network architecture and function largely remains missing [36].

Here, we address this gap by reconstituting cell-like vimentin networks in vitro using purified vimentin and clarified cell extracts. We use this system to connect spatial vimentin organization and mechanics in a physiologically relevant context and establish cytosolic control as a key regulator of vimentin network architecture. We show that: (i) vimentin forms complex, cell-like networks in the presence of cytosolic factors; (ii) network architecture and mechanics are dominated by extract composition rather than filament length or vimentin concentration; and (iii) extracts from different cell types generate distinct architectures, consistent with a cell-type-specific regulation of intermediate filament networks. Together, our results provide a conceptual and experimental bridge between well-established in vitro models on the single filament level and the complex intracellular environment, establishing a route to interrogate intermediate filament networks in a physiologically relevant context.

## Results

### Vimentin filaments form cell-like networks in cell extract

In cells, vimentin filaments form well-structured networks composed of fine meshes and bundles. For example, in mouse embryonic fibroblasts (MEFs), vimentin typically organizes into a dense, cage-like structure surrounding the nucleus and extending toward the cell periphery (Fig. 1A). Such organization reflects extensive crosslinking and bundling, which are hallmarks of vimentin network architecture [6], [14], [37], [38].

**Figure 1:**
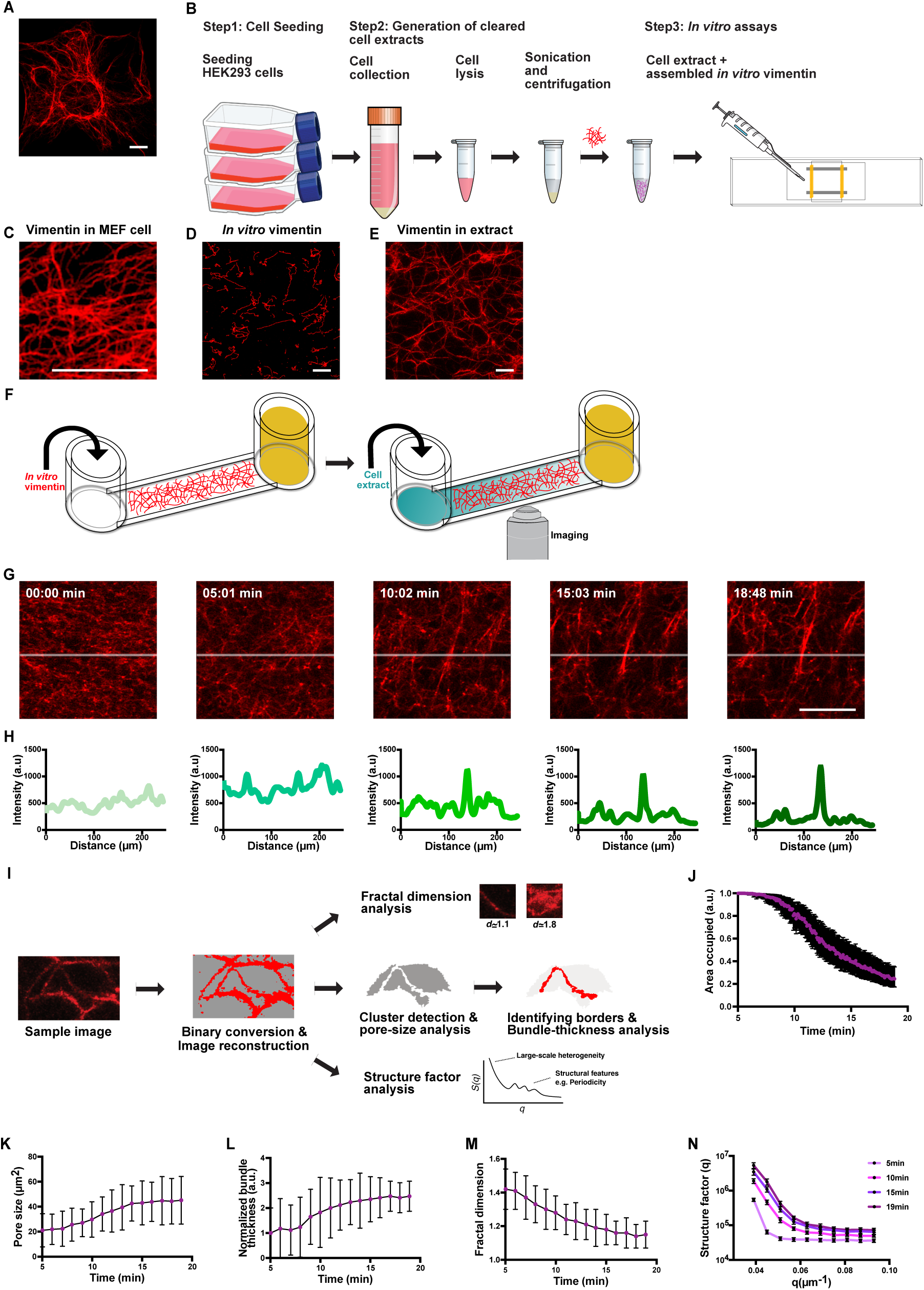
Vimentin forms cell-like networks in the extract. **(A)** Immunostaining of vimentin in a mouse embryonic fibroblast (MEF) cell. **(B)** Schematic representation of the extract-based pipeline used in this study. In step 1, HEK293 cells are seeded in flasks and grown to 80% confluence. In step 2, HEK293 cells are lysed and centrifuged to obtain clarified extracts. In step 3, assembled *in vitro* vimentin filaments are added to the clarified extracts for *in vitro* reconstitution assays. **(C)** Zoomed-in image of vimentin filaments in a MEF cell. **(D)** *In vitro* assembled vimentin filaments labeled with ATTO 647N on a glass coverslip. **(E)** *In vitro* vimentin forming a cell-like network in the extract. **(F)** Schematic representation of the experimental setup. *In vitro* assembled vimentin filaments were first added to one of the reservoirs of a channel on a glass coverslip. To prevent flow, the second reservoir was then closed. Cell extract was added to the first reservoir and slowly diffused into the channel. **(G)** Time evolution of the vimentin network formation. **(H)** Intensity profiles of vimentin network formation extracted from horizontal line scans across the corresponding images in (G). **(I)** Schematic of the image analysis pipeline used in this study. **(J)** Plot showing the time evolution of the area fraction occupied by vimentin in (G). The error bars indicate the variation range of the area fraction for the admissible range of intensity thresholds. Additional replicates are shown in Fig. S1A. **(K,L,M,N)** Plots showing time evolution of the mean pore size, normalized bundle thickness, fractal dimension and structure factor of the vimentin network shown in (G). The error bars indicate the standard deviation across three replicates. Scale bar = 10 μm.

To investigate how vimentin filaments assemble into networks in a cell-like biochemical environment, we used a cell extract-based approach previously employed to reconstitute microtubule interactions with their regulatory proteins [39], [40]. This strategy preserves near-native biochemical conditions while maintaining the experimental accessibility of an in vitro system. We generated clarified extract by growing HEK293 cells to 80% confluence (Fig. 1B, step 1), collecting and lysing the cells, and centrifuging the homogenate to obtain the cleared supernatant (step 2). HEK293 cells were chosen because they grow rapidly and thus support high-yield extract preparation, have been successfully used in extract-based reconstitution pipelines [39], and express vimentin [41]. In parallel, we assembled Atto-647N-labeled vimentin filaments for 16 h from 0.2 mg/mL recombinantly expressed, purified protein [19], [42] and introduced these filaments into the extract within glass flow chambers for imaging (step 3).

A magnified view (Fig. 1C) of the intracellular vimentin network shown in Fig. 1A illustrates the high degree of crosslinking and the intricate mesh architecture characteristic of vimentin in cells. In contrast, purified vimentin filaments in buffer do not show visible interactions or form higher-order structures (Fig. 1D). Strikingly, upon addition to the cell extract, the same filaments rapidly form a highly crosslinked and stable, percolated filament mesh that spans tens of microns and exhibits micron-scale pore structure – features we refer to here as a cell-like vimentin network (Fig. 1E). Although the mesh size in extract is larger than in cells – likely due to inevitable dilution during extract preparation – the resulting architecture recapitulates the extended connectivity and bundling of intracellular vimentin networks. To our knowledge, this is the first demonstration that purified vimentin filaments can assemble into a cell-like network architecture outside cells.

To follow the temporal evolution of network formation, we established a simple diffusion-mixing assay. Pre-assembled vimentin filaments were introduced into a glass microchannel connected to two reservoirs (Fig. 1F, left). After blocking one reservoir, we added extract to the opposite reservoir and immediately imaged the channel center as the extract diffused into the filament solution (Fig. 1F, right). An example time series (Fig. 1G, suppl. movie 1) shows that filaments initially exhibit pronounced thermal fluctuations (0 min), begin to crosslink and immobilize after ∼5 min, form bundles and a more coherent mesh by ∼10 min, and reach a stable network architecture by ∼15 min. Line-scan intensity profiles across the field of view (Fig. 1H) reveal the emergence of sharper peaks over time, corresponding to bundling and network consolidation.

To quantify network architecture, we developed an image analysis pipeline (Fig 1I). Images were thresholded, binarized, and cleared by removing small, disconnected particles. From these processed images, we extracted key network architecture parameters: pore size, (normalized) bundle thickness, and the area fraction occupied by vimentin. We also measured the fractal dimension – a dimensionless metric reporting on spatial complexity – and the structure factor *S(q)*, a Fourier-space measure commonly used in soft-matter and crystallographic analysis [43], [44] that reports on large-scale heterogeneity (for small *q*) as well as finer structural features (for large *q*). Details of the analysis pipeline are described in the Methods.

Consistent with visual inspection, the area occupied by vimentin decreases over time as filaments reorganize into bundles and crosslinked meshes (Fig. 1J, see Fig. S1A for further examples), increasing the space between bundles. Accordingly, pore size and bundle thickness increase as the network opens up (Fig. 1K,L). In parallel, the fractal dimension decreases (Fig. 1M), indicating a transition from an initially diffuse filament distribution toward a more consolidated architecture. The structure factor increases, particularly at small to intermediate *q* (Fig. 1N), which correspond to real-space length scales of ∼15-25 µm – notably larger than the mesh size. This indicates that the most pronounced architectural changes occur at supramesh dimensions, revealing the formation of extended bundles and larger domains. We used GFP as a freely diffusing marker of extract arrival and found that the onset of network formation coincides with the sharp rise in GFP intensity (Fig. S1B,C). This indicates that network formation is rapid once cytosolic components arrive and is likely diffusion-limited in this assay.

Together, these results demonstrate that purified vimentin filaments can assemble into robust, cell-like networks in cell extract, that this assembly occurs on the timescale of minutes, and that the resulting architecture remains stable thereafter.

### Vimentin network architecture is cell type-dependent

Clarified extracts contain endogenous vimentin, even though larger structures such as organelles and large clusters of pre-existing filaments are removed during preparation. This raises the question whether the in vitro assembled vimentin filaments form networks independently, or whether network formation requires residual endogenous vimentin in the extract. To address this, we generated cell extract from RPE-1 cells expressing endogenously GFP-labeled vimentin [14]. As in HEK293 extract, the in vitro assembled filaments formed extensive networks (Fig. 2A, right). Endogenous vimentin, in contrast, appeared sparse and fragmented and co-localized with the in vitro assembled network (Fig. 2A, left), suggesting that endogenous vimentin is incorporated into, and supported by, the network formed by the added filaments, rather than driving network formation itself. We next prepared extract from vimentin -/- MEF cells. Again, the pre-assembled filaments formed robust networks that closely resembled those formed in HEK293 extract (Fig. 2B), demonstrating that endogenous vimentin is not required for network formation.

**Figure 2:**
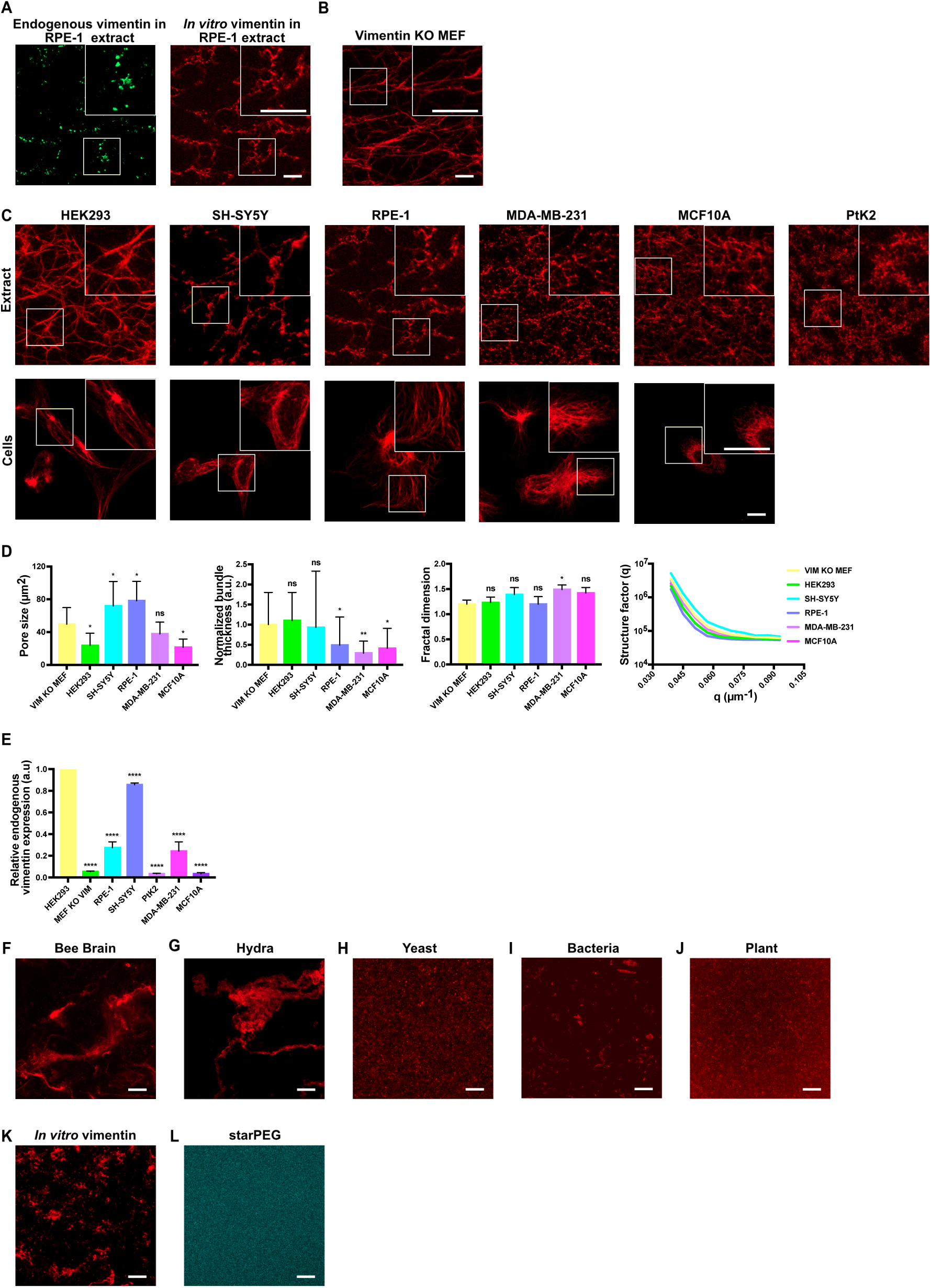
Vimentin network architecture is cell type-dependent. Representative images showing vimentin networks in **(A)** RPE-1 extract showing endogenously mEmerald-labeled vimentin (left) and *in vitro* assembled vimentin (right), **(B)** vimentin KO MEF extract. **(C)** Representative images showing vimentin networks in HEK293, SH-SY5Y, RPE-1, MDA-MB-231, MCF10A and PtK2 cell extracts, respectively (top). Lower panels show immunostaining of vimentin in the afore-mentioned cell lines. **(D)** Plots representing quantifications of pore size, normalized bundle thickness, fractal dimension and structure factor, respectively, for the above-mentioned cell extracts. **(E)** Quantification of immunoblot band intensities of endogenous vimentin in the cell lines from (C). Representative images showing vimentin networks in **(F)** bee brain extract**, (G)** hydra extract, **(H)** yeast extract, **(I)** bacterial extract, **(J)** plant extract and **(K,L)** in the presence of starPEG-(KA7)4 –TAMRA (**K**: vimentin channel, **L**: starPEG channel). Quantifications were done for three independent biological replicates (unpaired t-test, N = 3, mean ± SD, *P < 0.05, **P < 0.01, ***P < 0.001 and ****P < 0.0001); For **(D)** each data set was compared against its preceding data set, For **(E)**, all data sets were compared to HEK293. Scale bar = 10 μm.

To determine whether vimentin network architecture depends on the cell type from which the extract is derived, we prepared extract from a panel of mammalian cell lines representing diverse tissue origins. Fig. 2C shows networks formed from in vitro assembled filaments (top) alongside vimentin immunostaining in fixed cells (bottom), ordered by endogenous vimentin expression (Fig. 2E, Fig. S2G), from high (left) to undetectable (right). Strikingly, vimentin formed networks in every extract tested, including those from cell lines that express little to no endogenous vimentin. This indicates that the cytosolic factors required for network formation are present even in cells with minimal or no vimentin expression.

Despite this shared ability to support network formation, the detailed architectures varied substantially across extracts. Quantification (Fig. 2D) revealed no simple relationship between endogenous vimentin levels and network architecture. Extract from SH-SY5Y and RPE-1 cells produced the largest mesh sizes, whereas HEK293 and MCF10A extracts produced the smallest. Bundle thickness was highest in vimentin -/- MEF, HEK293, and SH-SY5Y extracts. In contrast, the fractal dimension and structure factor were similar across all cell types, ranging between ∼1.2 and ∼1.5, indicating moderately space-filling architectures with broadly conserved structural complexity.

These observations raise the question of whether the cytosolic factors that promote network formation are also present in species that, to current knowledge, do not express cytoplasmic intermediate filaments. To test this, we generated extract from bee brain, hydra, yeast, bacteria, and plant cells. Vimentin formed assemblies in bee brain and hydra extract, although these structures did not resemble typical vimentin networks (Fig. 2F,G). In contrast, yeast, bacterial, and plant extracts did not support any network-like structures (Fig. 2H-J). Whether insect intermediate filament-like proteins such as isomin from *Isotomurus maculatus* are bona fide intermediate filaments remains unclear [45], [46], and to our knowledge no intermediate filament-like proteins have been described in hydra. Overall, these results suggest that network formation depends on cytosolic factors present specifically in intermediate filament-expressing cell types.

To test whether generic cytosolic components could account for network formation, we modulated concentrations of Mg^2+^ and Ca^2+^ ions, which are commonly used to promote charge-based filament interactions [33], [34], [47]. Notably, purified filaments are already in 100 mM KCl, indicating that monovalent cations like K^+^ alone are insufficient to drive network formation. Consistent with previous reports, neither divalent cation induced cell-like networks; instead, both conditions produced local aggregates and clumps (Fig. S2A,B). We also tested starPEG, a positively charged multivalent synthetic polymer originally designed to mimic the electrostatic behavior of microtubule associated proteins [47]. Similar to divalent ions, starPEG was not able to reproduce cell-like vimentin networks (Fig. 2K,L).

These findings indicate that essential factors for network formation are biomolecules – most likely proteins – rather than generic cytosolic components such as simple ions. Consistent with this, heat-inactivated HEK293 extract failed to support network formation (Fig. S2C), instead producing images resembling those obtained from species lacking intermediate filaments. We note that, at least in our assay, controlled pre-assembly of filaments is required for network formation (Fig. S2E).

Taken together, our results show that vimentin network formation relies on cell type-specific cytosolic biomolecules and cannot be explained by vimentin expression alone or by nonspecific electrostatic interactions. The cytosolic environment therefore plays a guiding role in shaping vimentin network architecture.

### Vimentin network formation is robust, but its architecture is tunable

For a more detailed understanding of vimentin network architecture, we focused on HEK293 cell extract and systematically varied three key parameters: extract concentration (changing the availability of cytosolic network-organizing factors), vimentin concentration, and vimentin assembly time (shifting filament length, Fig. S3B,C), while keeping the other two parameters constant. Across perturbations, networks formed reproducibly, but their architectures varied. Increasing extract concentration (Fig. 3A) led to a strong reduction in pore size and a modest increase in bundle thickness. The fractal dimension remained nearly constant, whereas the structure factor increased markedly at low *q*, indicating large-scale density variations. These trends suggest that higher extract levels promote network compaction and long-range clustering (see Fig. S3A for further example images). Notably, network formation required a minimum of ∼ 1 mg/mL extract; below this threshold, filaments remained mobile and did not organize into networks.

**Figure 3:**
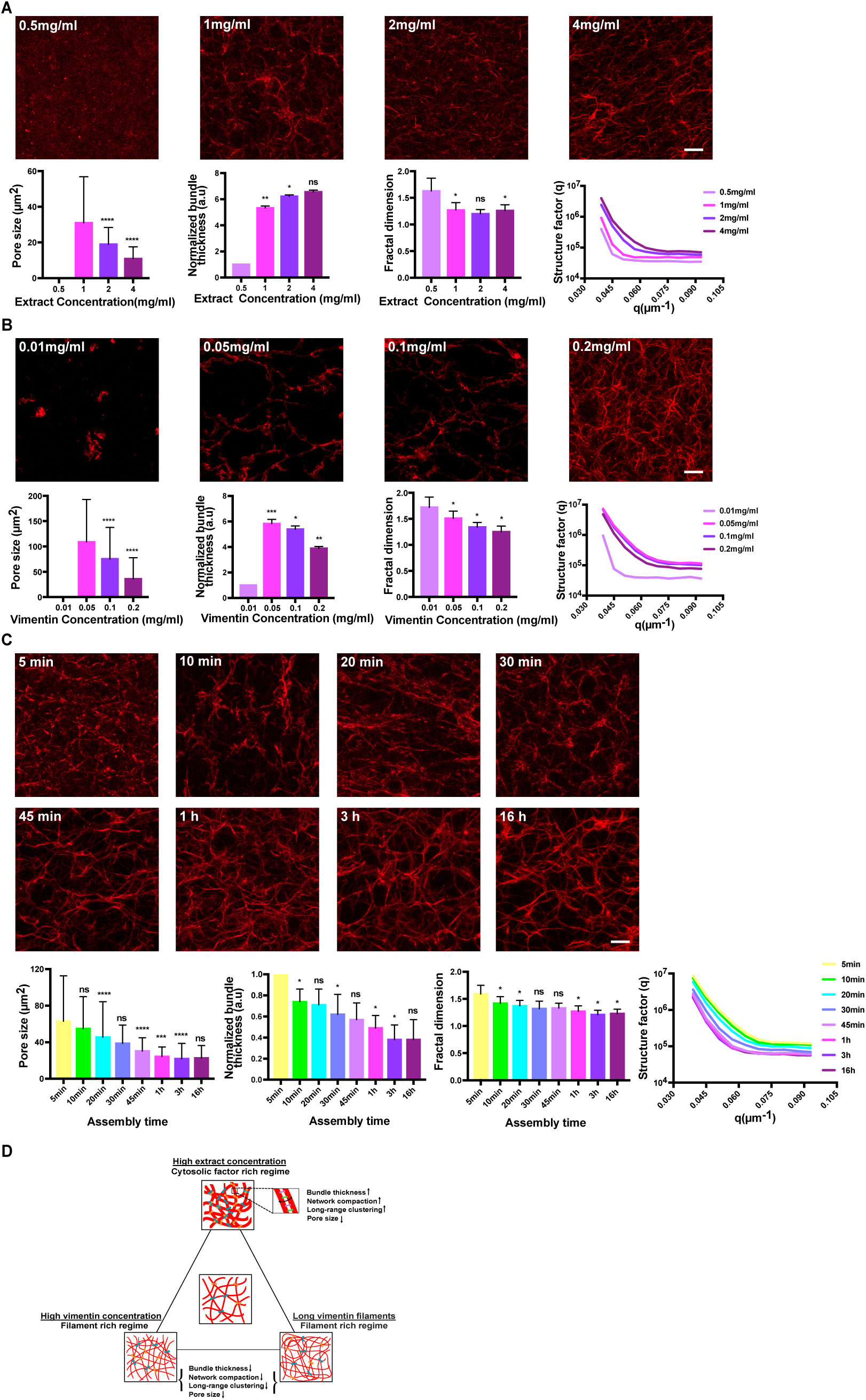
Vimentin network architecture depends on extract composition and filament properties. **(A)** Representative images showing vimentin networks in varying concentrations of cell extract (0.5 mg/mL, 1 mg/mL, 2 mg/mL, 4 mg/mL). Here, vimentin filaments were assembled at 0.2 mg/mL for 16 h. Lower panels show plots representing quantifications of pore size, normalized bundle thickness, fractal dimension and structure factor for varying extract concentrations. **(B)** Representative images showing vimentin networks with varying concentrations of *in vitro* assembled vimentin (0.01 mg/mL, 0.05 mg/mL, 0.1 mg/mL, 0.2 mg/mL). Here, vimentin filaments were assembled for 16 h and the extract concentration was kept at 1 mg/mL. Lower panels show plots representing quantifications of pore size, normalized bundle thickness, fractal dimension and structure factor for varying vimentin concentrations. **(C)** Representative images showing vimentin networks with varying assembly times (leading to varying filament lengths) of *in vitro* assembled vimentin (5 min, 10 min, 20 min, 30 min, 45 min, 1 h, 3 h, 16 h). Here, vimentin filaments were assembled at 0.2 mg/mL and the extract concentration was kept at 1 mg/mL. Lower panels show plots representing quantifications of pore size, normalized bundle thickness, fractal dimension and structure factor for varying assembly times (and thus filament lengths) of *in vitro* assembled vimentin. Values for bundle thickness were normalized to the smallest condition (that exhibit bundles within the network). Quantifications were done for three independent biological replicates (unpaired t-test, N = 3, mean ± SD, *P < 0.05, **P < 0.01, ***P < 0.001 and ****P < 0.0001); Here, each data set was compared against its preceding data set. **(D)** Schematic showing the vimentin network architectural changes between the filament rich regime and cytosolic factor rich regime. Scale bar = 10 μm. Images were contrast-adjusted for better visualization.

Increasing vimentin concentration (Fig. 3B) also reduced pore size, but produced thinner bundles. The fractal dimension decreased moderately and the structure factor remained almost unchanged. Thus, higher filament density yields finer and more homogeneous, less bundled networks. A minimal vimentin concentration of ∼ 0.05 mg/mL was required for network formation.

Increasing filament length (Fig. 3C, Fig. S3B,C) decreased pore size and bundle thickness and caused slight reductions in fractal dimension and structure factor. Longer filaments therefore generated denser, more uniformly connected networks without increasing bundling.

Together, these results support a framework in which network architecture reflects the balance between the availability of filaments on the one hand and cytosolic factors that structure the filaments into networks on the other (Fig. 3D). At high extract concentration, cytosolic factors are abundant and filaments become limiting: filaments are drawn together into denser meshes and slightly thicker bundles, producing small pores and long-range clustering. Consistent with this compaction, the vertical range occupied by the network in the flow chamber decreases at higher extract concentration, whereas filament concentration and length have no visible effect (Fig. S3D-F). At high vimentin concentration, many filaments compete for a fixed pool of cytosolic factors, pushing the system toward a filament-dominated regime. Here, bundling is reduced and the architecture becomes more homogeneous; however, the increased number of filaments still enhances overall connectivity, which can stabilize the network even in the absence of strong bundling. Longer filaments increase connectivity across larger distances without requiring bundling, generating dense yet more homogeneous networks.

### Cytosolic factors dominate vimentin network mechanics

The architectural differences that arise when varying extract concentration, vimentin concentration, or filament length suggest that these networks may also differ mechanically. To assess network mechanics, we monitored (i) the extent of network fluctuations, reasoning that mechanically stable networks should fluctuate less, and (ii) the mean-squared displacement (MSD) of the network’s center of mass (based on the imaged field of view). We quantified fluctuations as the fraction of network area that changed between two consecutive images (0% meaning perfect overlap, and 100% meaning no overlap). For the MSD, we extracted the anomalous exponent ⍺ from the scaling relation MSD(Δt) ∝ Δt^⍺^; freely floating filaments should yield ⍺ ≈ 1, whereas 0 < ⍺ < 1 reflects constrained (subdiffusive) motion.

We first compared networks assembled at the extremes of the experimentally accessible range: 1 mg/mL, the minimum required for visible network formation, and 4 mg/mL, the maximum achievable. Heatmaps generated from overlaying binary images across the entire observation period (Fig. 4A) reveal that the networks at 1 mg/mL exhibit large spatial fluctuations, whereas those at 4 mg/mL are almost static. Quantification confirms this: relative fluctuations exceed 15% at 1 mg/mL but fall below 3% at 4 mg/mL (Fig. 4B). MSD analysis yields an anomalous exponent close to diffusive behavior at low extract (⍺ = 0.95 ± 0.01) but strongly subdiffusive at high extract (⍺ = 0.22 ± 0.12) (Fig. 4C, Fig. S4A). Thus, a fourfold increase in extract concentration generates qualitatively different mechanical behavior.

**Figure 4:**
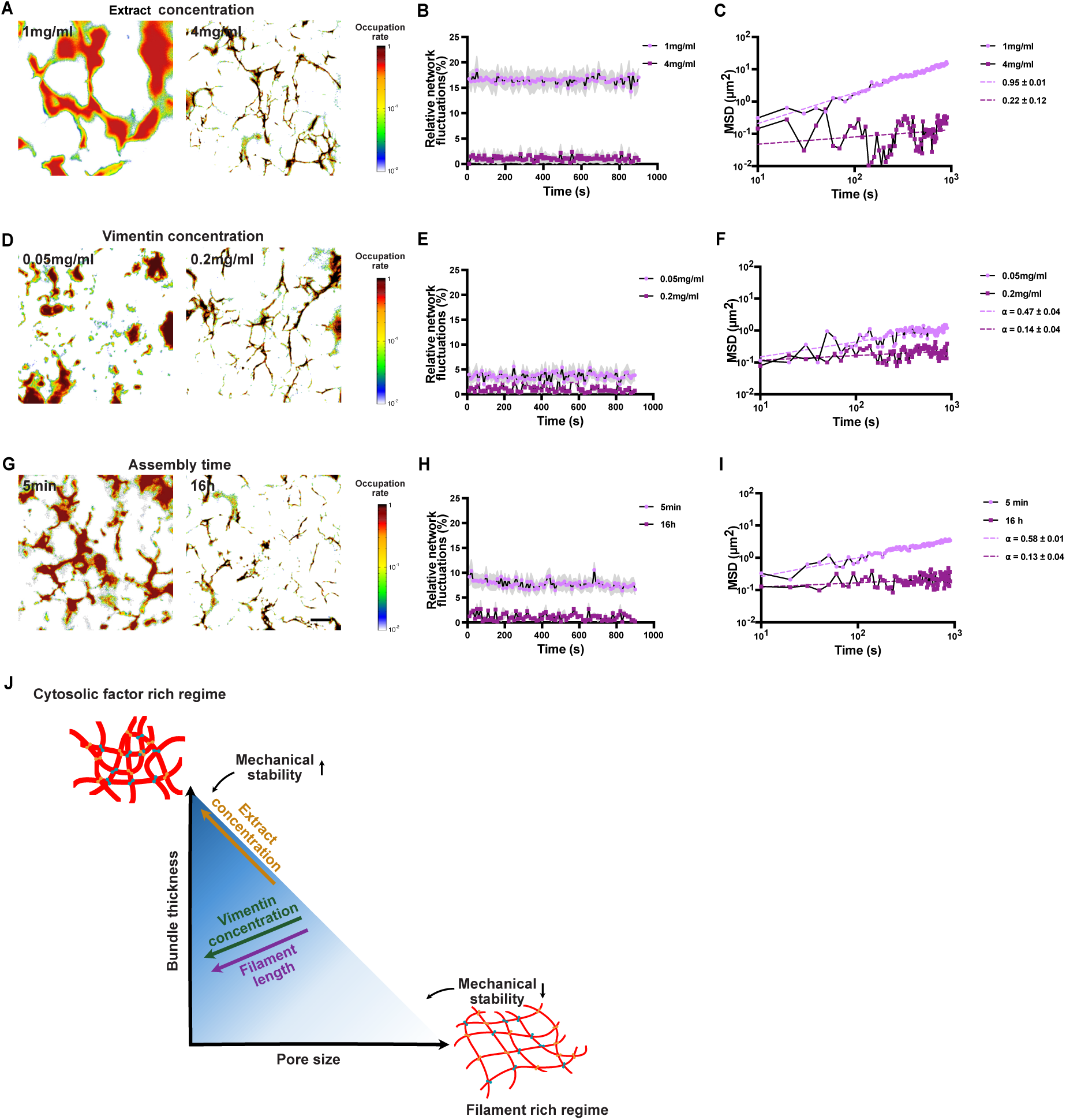
Extract concentration is the main determinant of network mechanics. **(A)** Representative heatmaps showing relative structural variations, measured as occupation rates, of the vimentin network over a period of 15 min with varying concentrations of cell extract (1 mg/mL, 4 mg/mL). Here, vimentin filaments were assembled at 0.2 mg/mL for 16 h. **(B)** Plot showing relative network fluctuations with varying extract concentrations. **(C)** Plot showing the mean-squared displacement (MSD) of the centre of mass of vimentin networks with varying extract concentrations. **(D)** Representative heatmaps showing relative structural variations of the vimentin network over a period of 15 min with varying concentrations of *in vitro* assembled vimentin (0.05 mg/mL, 0.2 mg/mL). Here, vimentin filaments were assembled for 16 h and the extract concentration was kept at 4 mg/mL. **(E)** Plot showing relative network fluctuations with varying *in vitro* vimentin concentrations. **(F)** Plot showing the MSD of the centre of mass of vimentin networks with varying *in vitro* vimentin concentrations. **(G)** Representative heatmaps showing relative structural variations of the vimentin network over a period of 15 min with varying assembly time (and thus filament lengths) of *in vitro* assembled vimentin (5 min, 16 h). Here, vimentin filaments were assembled at 0.2 mg/mL and extract concentration was kept at 4 mg/mL. **(H)** Plot showing relative network fluctuations with varying assembly time of *in vitro* assembled vimentin. **(I)** Plot showing the MSD of the centre of mass of the vimentin network fluctuations with varying assembly time of the *in vitro* assembled vimentin. The shaded regions in B,E,H show the standard deviation; data shown is pooled from three independent replicates. Additional replicates of MSD analysis are shown in Fig. S4. The centre of mass rearrangements follow an anomalous diffusion MSD(Δt)∝Δt^α^. Here, α is the anomalous exponent, which reflects the overall dynamics of the network: sub-diffusion (slower than diffusion) for 0<α<1, normal diffusion for α=1, and super-diffusion (faster than diffusion) for 1<α<2; For a detailed explanation on the quantification of relative network fluctuations and MSD, refer to the methods section. Scale bar = 10 µm. (**J**) Schematic summary linking network architecture to mechanical stability. Network mechanics arise from the interplay between filament supply and cytosolic network-organizing factors. Extract concentration exerts the strongest control, simultaneously decreasing pore size and increasing bundle thickness, which together produce highly stable networks. In contrast, increasing vimentin concentration or filament length reduces pore size but also decreases bundle thickness, generating competing effects.

Next, we varied vimentin concentration (0.05 mg/mL and 0.2 mg/mL) while keeping extract concentration high. Although these networks differ noticeably in density (Figs. 3B, 4D), their mechanical behavior is surprisingly similar. Both exhibit low fluctuations (< 5%, Fig. 4E) and subdiffusive MSDs (⍺ = 0.47 ± 0.04 and ⍺ = 0.14 ± 0.04) (Fig. 4F, Fig. S4B). Thus, increasing filament abundance makes networks more stable, but the effect is moderate compared to changes in extract concentration.

Finally, we varied filament assembly time to obtain different length distributions at constant vimentin and extract concentrations. Networks composed of shorter filaments (5 min assembly) show increased fluctuations (Fig. 4G,H) and higher MSDs (Fig. 4I, Fig. S4C), but these differences are again modest relative to those induced by extract concentration.

Overall, extract concentration – and therefore the availability of cytosolic network-organizing factors – has the strongest impact on vimentin network mechanics. Integrating these data with the architectural analysis (Fig. 3) suggests the following view (Fig. 4J): Pore size is the architectural parameter that varies most strongly across conditions. Smaller pores generally correspond to greater connectivity and / or steric hindrance. Given the very low persistence length of vimentin filaments (∼ 1 µm), mechanical stability likely arises from physical cross-connections between filaments rather than steric entanglement. Bundle thickness also contributes: bundles are mechanically stiffer than individual filaments. Extract concentration affects both pore size (decreasing it) and bundle thickness (increasing it), producing much more stable networks. In contrast, changes in vimentin concentration or filament length produce opposing effects: pore size decreases (stabilizing), but bundle thickness decreases as well (destabilizing). This compensation likely explains why these parameters exert only modest influence on mechanics overall.

## Discussion

Our observations support a model in which vimentin network architecture is determined by the balance between filament availability and cytosolic network-organizing factors. Cytosolic factors act as crosslinkers and bundling agents: when abundant, they draw filaments into compact, bundled, and heterogeneous architectures, whereas when scarce, filaments distribute more uniformly and connectivity arises from filament abundance and length rather than bundling. Network formation requires heat-sensitive, cell type-specific biomolecules present in extracts from mammalian cells that express cytosolic intermediate filaments – even when they do not express vimentin itself – and is lost in heat-inactivated extract or extracts from species lacking cytosolic intermediate filaments. Thus, network assembly requires specific biomolecules rather than generic charge-based interactions: although vimentin is a strong polyelectrolyte [34], [48], mono- and divalent cations induce only non-specific aggregation and do not generate extended, percolated, cell-like network architectures. Together, these observations place constraints on the nature of the organizing factors and suggest that one or more proteinaceous components capable of multivalent filament interactions act as effective linkers and organizers of vimentin filaments. Networks assembled from identical biochemical components nonetheless adopt architectures depending on the length of filaments at the onset of network formation, consistent with observations on crosslinked actin networks in which assembly kinetics and dynamic arrest shape final architectures [44], [49].

In cells, vimentin networks are additionally shaped by anchoring to the nuclear envelope, membranes, and other intracellular structures, which likely contribute to characteristic features such as the perinuclear cage [50]. These boundary conditions are necessarily absent in clarified extracts. The ability of vimentin to form extended, mechanically stable networks in extract demonstrates that anchoring is not required for network formation per se, but likely acts as an additional layer of spatial control.

While vimentin organization in cells is influenced by microtubules [14], [51], [52], our assays demonstrate that network formation can occur in their absence, indicating that microtubules are not strictly required for network formation under these conditions (Fig. S2F). Because some of the cell lines used here are epithelial, a potential role for keratins in vimentin network formation may be considered. However, robust vimentin networks also form in extract from MEFs, which display a mesenchymal phenotype and are not dominated by keratin, indicating that keratins are not required for network assembly. Moreover, keratin expression is low in some epithelial cell types such as MCF10A cells [53], yet their extracts still support vimentin network formation. While keratins and other intermediate filament types may modulate network architecture, our results demonstrate that vimentin network assembly in extract does not require keratin.

Post-translational modifications (PTMs) of vimentin are known to regulate filament dynamics, interactions, and mechanics in cells [54], [55]. In our system, vimentin networks form from recombinant vimentin expressed in bacteria, which is expected to lack eukaryotic PTMs, and network assembly does not require endogenous vimentin. While recombinant filaments could in principle acquire modifications upon exposure to cytosolic enzymes, network formation occurs rapidly at the timescale of seconds to minutes and does not undergo further visible remodeling thereafter, arguing against a dominant role for (slow) enzymatic modification in driving assembly. PTMs therefore play likely modulatory rather than essential roles in network formation under these conditions.

Differences in network architecture across extracts suggest that multiple factors may contribute, potentially in cell-specific combinations, which could explain both cell-specific differences in vimentin organization and function and how vimentin network architecture differs from that of other intermediate filaments. Together, these findings reveal that vimentin networks self-organize through cell-type–specific cytosolic factors acting on a filament pool with tunable physical properties. These results also raise intriguing physiological implications: many disease states – including fibrosis, cancer progression, and inflammatory activation – are associated with drastic remodeling of vimentin networks [56], [57], [58], [59], [60]. Our results point to the possibility that not only vimentin expression, but also the expression or activity of cytosolic factors controlling network architecture may change in these contexts. Identifying these factors now emerges as a defined mechanistic question, and our extract-based reconstitution provides a tractable route to address it. Uncovering vimentin-associated factors and how their expression relates to disease states may ultimately provide mechanistic insight and potential therapeutic strategies.

## Acknowledgements

We thank Gijsje Koenderink for critically reading the manuscript and providing valuable feedback. We are grateful to Manuel Théry, Barbara Niemeyer, John Eriksson, Gaudenz Danuser, and Franziska Lautenschläger for generously providing mammalian cell lines, to Stefan Diez for providing StarPEG, and to Albrecht Ott, Mariana Romeiro Motta, Uli Müller, and Bruce Morgan for providing non-mammalian cells and tissues. We thank Sarah Köster and Susanne Bauch for technical assistance with vimentin purification. D.S., B.K. and L.S. were supported by a grant from the European Research Council (ERC Starting Grant, grant agreement no. 101115795) awarded to L.S. C.J. was supported by the Institut Curie, the French National Research Agency (ANR) award ANR-20-CE13-0011, and the Fondation pour la Recherche Medicale (FRM) grants DEQ20170336756 and MND202003011485. A.S.J. was supported by the European Union’s Horizon 2020 research and innovation program under the Marie Skłodowska-Curie grant agreement no. 675737, and the FRM grant FDT201904008210.

## Author contributions

Conceptualization, L.S.; funding acquisition, L.S.; methodology, D.S., R.S., M.G., B.K., A.S.J., C.J.; investigation, D.S.; analysis, D.S., R.S., L.S.; supervision, L.S.; writing, L.S., D.S.

## Declaration of generative AI and AI-assisted technologies in the writing process

During the preparation of this work the authors used ChatGPT-5.2 (OpenAI) for language editing and stylistic refinement. After using this tool, the authors reviewed and edited the content as needed and take full responsibility for the content of the published article.

## Materials and methods

### Vimentin purification, labeling and assembly

Vimentin C328NGGC was recombinantly expressed [16], [42], [61] and stored at −80 °C in a buffer containing 1 mM EDTA (ethylenediamine tetraacetic acid), 0.1 mM EGTA (ethylene glycol-bis (β-aminoethyl ether) -N,N,Nʹ,Nʹ-tetraacetic acid), 8 M urea, 0.1–0.4 M KCl, 1 mM DTT (dithiothreitol) and 5 mM Tris (tris (hydroxymethyl) aminomethane) at pH 7.5 [16]. After thawing, monomers were fluorescently labeled with ATTO647N dye (AD 647N-41, AttoTech, Siegen, Germany) as described in refs [19], [62], [63]. Labeled and unlabeled monomers were then combined such that 8% of the total monomer were fluorescently labelled.

Tetramers were reconstituted by dialyzing the protein first against 6 M urea in 50 mM phosphate buffer (PB) pH 7.5, then sequentially against decreasing urea concentrations (4, 2, and 0 M) in 2 mM phosphate buffer (pH 7.5). This was followed by a final overnight dialysis at 4 °C in urea-free 2 mM phosphate buffer. Filament assembly was initiated by transferring the tetramers into assembly buffer consisting of 100 mM KCl and 2 mM phosphate buffer at pH 7.5, followed by incubation at 37 °C overnight or for the indicated duration [16], [64].

### Cell culture

Human Embryonic Kidney cells (HEK293, DSMZ, ACC305) were cultured in DMEM media (Fisher Scientific, 42430025) supplemented with heat inactivated 10% Foetal Bovine Serum (FBS Gibco A5670701) and 1% penicillin-streptomycin solution (Gibco, 15070063). Wild Type WT Mouse Embryonic Fibroblasts (MEFs) with vimentin and vimentin knockout (Vim KO)

MEFs were a kind gift from John Eriksson (Abo Åkademi University, Turku, Finland). These cells were cultured in DMEM media (Fisher Scientific, 42430025), supplemented with heat inactivated 10% FBS (Gibco A5670701) and 1% penicillin-streptomycin solution (Gibco, 15070063). SH-SY5Y (human neuroblastoma cell line) cells were a kind gift from Barbara Niemeyer (Saarland University, Germany). SH-SY5Y and Male potoroo kidney epithelial cells (PtK2) cells were cultured in DMEM/F12 (Gibco, 31331028) media supplemented with 10% FBS (Gibco, A5670701) and 1% penicillin-streptomycin solution (Gibco, 15070063). Immortalized human Retinal Pigment Epithelial cells (hTERT-RPE-1) endonegously expressing mEmerald-vimentin and mTagRFPT-tubulin as described in [14] were a kind gift from Gaudenz Danuser. These cells were cultured in DMEM/F12 (Gibco31331028) media supplemented with 10% FBS (Gibco A5670701), 1% GlutaMAX (35050061, Gibco) and 1% penicillin-streptomycin solution (Gibco, 15070063). MDA-MB-231 (triple-negative human breast cancer cell line) and MCF10A (non-tumorigenic human mammary epithelial cell line) cells were a kind gift from Franziska Lautenschlager (Saarland University, Germany). MDA-MB-231 cells were cultured in DMEM/F12 (Gibco, 31331028) media supplemented with 10% FBS (Gibco A5670701), 1% GlutaMAX (Gibco, 35050061) and 1% penicillin-streptomycin solution (Gibco, 15070063). MCF10A cells were cultured in DMEM/F12 (Gibco, 31331028) media supplemented 5% horse serum (Thermo Fisher Scientific, #16050-122) 20 ng/ml Epidermal Growth Factor (EGF) (PeproTech, 100-15), 0.5 µg/ml hydrocortisone (Sigma-Aldrich, H-0888), 100 ng/ml cholera toxin (Sigma-Aldrich, C-8052), 10 µg/ml insulin (Sigma-Aldrich, I9278) 10 mM HEPES and 1 % penicillin-streptomycin solution (Gibco, 15070063). All cell lines were maintained at 37 °C in a humidified incubator with 5% CO2.

### Preparation of cell extract

Mammalian cell extract: Two million HEK293 were seeded on T175 flasks. Cells were grown till ∼80% confluency. After removal of medium, cells were washed with 1X PBS. Cells were detached by adding 2 mL of Trypsin. The cells were collected with 6 mL of complete DMEM medium and transferred to a tube for centrifugation (450 g, 4°C, 10 min). The supernatant was removed and the cell pellet was re-suspended in 200-300 µl of lysis buffer containing BRB80 buffer (80 mM K-Pipes pH 6.8, 1 mM EGTA and 1 mM MgCl2), containing 0.05% Triton X-100 and protease inhibitors (10 µg ml^−1^ leupeptin, aprotinin and 4-(2-aminoethyl) benzenesulfonyl fluoride; Sigma-Aldrich). All the steps after the collection of cells were done on ice. The cell suspension was transferred to a 1.5 mL Beckman ultracentrifuge tube. The cells were lysed by pipetting up and down on ice for 5-7 mins and sonicated with three short pulses using the MS-72 probe (Bandelin Sonoplus), amplitude 12%. The sample was ultra centrifuged in the Beckman Ti70 rotor at 33,800 g, 4°C for 30 min in the Beckman Coulter Optima MAX-XP ultracentrifuge. The supernatants were aliquoted to 10-20 µl in cryovials, snap frozen in liquid nitrogen and stored at -80°C for upto three weeks [39].

Bee brain extract: Dissected brains from the bees (*Apis mellifera*) were manually crushed in BRB80 buffer to get the crude extract. Further, the above-mentioned protocol for mammalian cell extract was followed to prepare the clarified bee brain extract.

Hydra extract: Whole hydra (*Hydra vulgaris*) were manually crushed in BRB80 to get the crude extract. Further, the above-mentioned protocol for mammalian cell extract was followed to prepare the clarified hydra extract.

Bacterial extract: Cells were cultured by inoculating 3 mL of autoclaved LB medium with 5 µL of frozen cell stock carrying the plasmid pBest-OR2-OR1-Pr-UTR1-T500. This inoculation step served only to initiate growth and did not involve induction. Cultures were incubated overnight at 37 °C with shaking at 130 rpm until fully saturated. The following day, 1.5 mL of culture was transferred into a 2 mL microcentrifuge tube and centrifuged at 5,000 × g for 5 min at 4 °C. The supernatant was carefully discarded, and the pelleted cells were briefly centrifuged again at the same speed for 1 min to remove residual medium. Any remaining liquid was removed with a pipette without disturbing the pellet. Cell pellets were resuspended in lysis buffer containing 20 mM Tris-Cl (pH 7.5), 150 mM NaCl and 2 mM ATP. All handling steps were performed on ice. Cell lysis was performed using a Mini-Beadbeater with approximately 200 µL of 0.1 mm silica beads. Samples were subjected to four 30-second bead-beating cycles, with a 1-minute cooling period on ice between cycles to prevent heat buildup. Extract was clarified by centrifugation at 12,000 rpm for 20 min at 4 °C. The resulting supernatant was collected and stored at -80°C.

Yeast extract: Yeast extract was prepared as described in [65].

Plant extract: Fresh plant material (*Arabidopsis thaliana*) was ground to a fine powder in liquid nitrogen. The frozen powder was transferred to a pre-chilled tube and resuspended in 1 mL of extraction buffer (50 mM Tris pH 7.5, 150 mM NaCl, 10% glycerol, 2 mM EDTA, 5 mM DTT, 1% Triton X-100, 10 µl/mL plant protease inhibitor (Sigma #P9599), 20 µl/mL Phosphatase Inhibitor Cocktail (Sigma #P5726, #P0044)). Samples were incubated on ice for 30 minutes to allow efficient solubilization. The extract was clarified by centrifugation at 10,000×g for 5 minutes at 4 °C and the supernatant was transferred to a new tube. To ensure removal of remaining debris, the supernatant was centrifuged again at maximum speed. This clarification step was repeated until no visible pellet remained and the final clarified supernatant was collected and stored at -80°C.

### Cleaning of cover glasses

Cover-glasses were wiped with lint-free KimWipe^TM^ (Kimberly-Clark Professional™ 33670-04) tissues and 70% ethanol, incubated in acetone for 30 min, followed by 96% ethanol for 15 min (gentle agitation at room temperature). The cover-glasses were further rinsed in ultrapure water, incubated in Hellmanex III solution (2% in water, Hellmanex) for 2 h (gentle agitation at room temperature), washed in ultrapure water and air dried.

### *In vitro* reconstitution assays with cell extract

*In vitro* reconstitution assays were performed in 20 μl flow chambers that were made from sandwiching two pieces of cover-glasses using two strips of double-sided adhesive tape (70 μm height, 0000P70PC3003, LiMA, Couzeix, France). The assembled vimentin filaments were added to the cell extracts at a concentration of 0.04 mg/mL. The chamber was then sealed with Valap (a mixture of vaseline, lanolin and paraffin in equal parts) before imaging. For time evolution of vimentin networks in Fig. 1G, the chamber slides (ibidi, 80608) were stuck onto the cover glasses and secured onto the stage of the objective.

### Immunofluorescence

∼ 0.2 × 10^6^ cells/well of a 6-well plate were seeded onto glass coverslips. Cells were washed with 1x PBS and fixed with ice cold methanol for 5 min on ice, washed thrice in 1x PBS (3 min each). Cells were blocked in 3% BSA + 0.1% Tween prepared in 1X PBS, for 20 min. Incubation with vimentin primary antibody (D21H3 CST, 5741), dilution 1:200, was performed for 60 min at RT and with secondary antibody (Invitrogen, A-21245) for 30 min at RT, with washes in between using 1× PBS. Cells were washed thrice in 1x PBST after secondary antibody incubation. Further the cells were mounted using Fluromount-G (Invitrogen).

### Imaging

Fixed cells and *in vitro* samples (including time-evolution of vimentin network formation) were imaged on a Zeiss LSM900 confocal microscope with Airyscan equipped with an Axiocam 705 Mono camera using a 63x oil immersion objective (Immersol 518 F, NA=1.4; Plan Apochromat; WD = 0.19 mm; NA 1.4) at 1.3x digital zoom, Laser module 5, URGB (Laser line 640 nm was used). Confocal z-stacks were acquired at intervals of 2 μm for vimentin networks and at 0.5 μm for imaging fixed cells. Images were acquired using ZenBlue software (version 3.2, Zeiss).

Timelapse videos for vimentin network fluctuations were imaged using Nikon ECLIPSE Ti2 epifluorescence microscope. Videos were acquired with a 300 ms exposure (Light source - Epi-FL module with LED 640 nm coupled with a motorized epi filter turret) using a 60x oil objective (Nikon Plan Apo, NA = 1.42, WD = 0.15 mm).

### Protein quantification

The total protein concentration of the cell extract was determined using Pierce detergent compatible Bradford assay kit (Thermo Fischer Scientific), following the manufacturer’s instructions. The sample to reagent ratio was 1:30, was incubated at room temperature for 10 min and absorbance was measured at 595 nm with a plate reader. Absorbance values of BSA standard dilutions were plotted and fitted to a linear equation, and total protein concentrations of the cell extract were obtained.

### Western blotting

The samples were denatured for 5 min with 4x Laemmli buffer (Bio-Rad). 20 μg of the cell extract was loaded onto a 10% SDS-PAGE gel. Resolved proteins were transferred onto nitrocellulose membrane. Immunoblots were blocked using 5% non-fat milk prepared in 1x TBS. Immunodetection: primary antibody against vimentin (D21H3 CST, 5741), 1:1000; GAPDH (Proteintech, 60004-1-Ig), 1:2000; both overnight incubation at 4 °C, secondary antibody (IRDye800CW, IRDye680RD), 1:15000 at room temperature for 1 h. Blots were rinsed thrice with 1x TBST in between incubation. The blots were imaged via ChemiDoc using optimal exposures.

### Image analysis and quantification

All images and videos were contrast adjusted for better visualization in Fiji [66]. Line profiles in Fig. 1H and band intensities for western blot in Fig. 2E were analyzed in Fiji [66].

The methodology for extracting the structural characteristics of the vimentin network is outlined below. We begin by detailing the image refinement steps necessary to reconstruct each dataset in a manner that faithfully preserves the features of the original experimental images. Following this, we describe the analytical procedures used to quantify key structural features, including fractal dimension, structure factor, pore size, bundle thickness, and network fluctuations.

Reconstruction of images: Grayscale images were first converted into 2D data arrays in MATLAB. To isolate the vimentin network, a global intensity threshold was applied, converting the grayscale image into a binary one. As the threshold increases, the image transitions from a densely connected structure to one where discrete filament become visible, reminiscent of a percolation transition. To investigate whether a critical threshold exists, we varied the intensity threshold and measured both the fraction of occupied sites and the size of the largest connected cluster. The occupied fraction decreases smoothly with a varying behaviour between different images, with no clear threshold indicating a sharp transition. Next, we applied the Hoshen-Kopelman algorithm to detect clusters [67], assuming connectivity via nearest neighbours. We also tested an extended version including next-nearest neighbours, which yielded qualitatively similar results. We then tracked the size of the largest cluster as a function of intensity threshold, which also exhibited a smooth decay. From the combined behaviour of the occupied fraction and the largest cluster size, we manually selected the optimal intensity threshold for each image.

Fractal dimension: To quantify the spatial complexity of each vimentin network image, we computed the fractal dimension using a box-counting approach [43]. This method captures not only how much of the image is occupied by filaments but also how that occupation is spatially organized. In this context, a thin, linear filament yields a fractal dimension close to 1, whereas a densely packed region of vimentin approaches a dimension of 2. The procedure involves overlaying a grid of square boxes of size ε onto the binary image and counting the number of boxes N(ε) that contain at least one occupied pixel. This counting is repeated for progressively smaller ε, and the fractal dimension d is determined as the slope of the linear fit to the log-log plot of N(ε) versus 1/ε:

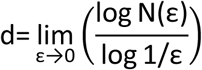

This scaling-based method effectively distinguishes between compact clusters - where N(ε) grows slowly with decreasing ε, resulting in higher d - and sparse or elongated structures, which exhibit faster growth in N(ε) and thus lower d.

Structure factor: To extract additional structural information from the vimentin network, we computed the static structure factor S(q), a method analogous to those used in solid state physics. This approach transforms spatial patterns into reciprocal space, enabling the identification of characteristic length scales and degrees of spatial order [68]. Starting from the binary image of the network, we compute the two-dimensional Fourier transform of the spatial density field. The structure factor S(q) is then obtained by radially averaging the squared magnitude of the Fourier-transformed field:

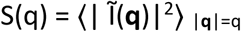

where Ĩ(**q**) is the Fourier transform of the intensity field Ĩ(**r**), and q=|**q**| denotes the spatial frequency. Extrema in the S(q) curve indicate dominant structural features such as inter-filament spacing or characteristic pore sizes, and the low-*q* behaviour reflects large-scale heterogeneity or clustering.

Pore size analysis: To quantify the pore structure within the vimentin network, we first adjusted the intensity threshold for each image to produce a binary image in which filamentous regions formed a well-connected network as far as possible. The pores, defined as the void spaces enclosed by filaments, were then identified by applying the cluster detection algorithm to the inverted binary image, where pores correspond to connected regions of background pixels. Large open regions connected to the image borders were assumed to be part of the external background and were excluded manually to avoid overestimating pore sizes. For each processed image, we computed the distribution of pore areas and extracted the mean pore size as a representative structural metric.

Bundle thickness: We utilized both pore geometry and visual calibration to estimate the average thickness of filamentous bundles within the vimentin network. First, we manually identified the thinnest visible filament and assigned its width as the reference filament width. Using the binary images in which pores had been segmented, we measured the average width of the borders separating neighbouring pores. These borders correspond to the filament bundles. The mean bundle thickness was then calculated by dividing this average border width by the reference single filament width, yielding a dimensionless measure of relative bundle thickness. This approach provides a practical estimate of bundling in the network without requiring sub-pixel resolution or direct filament tracing.

Network structural fluctuations: By analyzing time-lapse image sequences, we quantified temporal fluctuations in the vimentin network structure by tracking the center-of-mass of the network in each frame. For a given binary image at time *t*, the center-of-mass position **r**c.m. was calculated based on the spatial distribution of filament locations, defined as

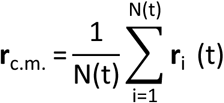

where **r**i(t) is the position vector of the i-th occupied pixel and N(t) is the total number of occupied pixels at time *t*. To characterize the dynamics of the structural fluctuations, we computed the mean squared displacement of the center-of-mass over time lags Δt as: MSD(Δt) = ⟨|**r**c.m.(t+Δt)−**r**c.m.(t)|^2^⟩t, where ⟨…⟩t denotes a time average over all valid initial time points t. For systems exhibiting diffusive-like fluctuations, the MSD scales linearly with time lag, and the effective diffusion coefficient D can be extracted from the relation MSD(Δt)=4DΔt in two dimensions. However, we observe that the center of mass rearrangements follow an anomalous diffusion MSD(Δt) ∝ Δt^α^. Here, α is the anomalous exponent [69], which reflects the overall dynamics of the network: sub-diffusion (slower than diffusion) for 0<α<1, normal diffusion for α=1, and super-diffusion (faster than diffusion) for 1<α<2. This analysis provides a coarse-grained measure of how the global network structure fluctuates over time, reflecting large-scale rearrangements or drift in the network morphology. As an alternative approach to quantify network fluctuations, we analyzed pairs of successive network images to assess their relative differences. For this purpose, we generated inverted binary images and counted the number of occupied pixels in consecutive time frames, denoted as N(t) and N(t+1), respectively. We also measured the number of pixels occupied in both frames, denoted as Noverlap(t). The relative structural variation of the network at time t was then calculated as 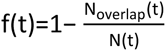. Additionally, we overlaid the binary image data over the entire observation period to construct heat map representing network fluctuations. In this representation, each element (i, j) of the 2D array is assigned an occupation probability 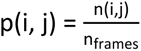 where nframes is the total number of analyzed frames and

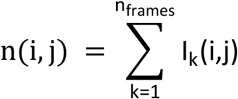

with Ik(i,j) being the binary intensity value at position (i, j) at time frame k. Thus, p(i, j) ranges within [0, 1], reflecting the fraction of time in which the (i, j) element has been a part of the network. By choosing a threshold probability pc(i, j) = 0.75 (or pc(i, j) = 0.5), we quantified the degree of network dynamics in each heat map using the ratio r = Nc / Ntot, where Nc is the number of pixels with occupation probability less than pc(i, j) and Ntot is number of pixels with non-zero occupation probability, respectively.

## References

[1] G. Dutour-Provenzano and S. Etienne-Manneville, ‘Intermediate filaments’, Curr. Biol. CB, vol. 31, no. 10, pp. R522–R529, May 2021, doi: 10.1016/j.cub.2021.04.011.

[2] I. Szeverenyi et al., ‘The Human Intermediate Filament Database: comprehensive information on a gene family involved in many human diseases’, Hum. Mutat., vol. 29, no. 3, pp. 351–360, Mar. 2008, doi: 10.1002/humu.20652.

[3] H. Herrmann and U. Aebi, ‘Intermediate Filaments: Structure and Assembly’, Cold Spring Harb. Perspect. Biol., vol. 8, no. 11, p. a018242, Nov. 2016, doi: 10.1101/cshperspect.a018242.

[4] C. Leduc and S. Etienne-Manneville, ‘Intermediate filaments in cell migration and invasion: the unusual suspects’, Curr. Opin. Cell Biol., vol. 32, pp. 102–112, Feb. 2015, doi: 10.1016/j.ceb.2015.01.005.

[5] E. Infante and S. Etienne-Manneville, ‘Intermediate filaments: Integration of cell mechanical properties during migration’, Front. Cell Dev. Biol., vol. 10, Aug. 2022, doi: 10.3389/fcell.2022.951816.

[6] M. G. Mendez, S.-I. Kojima, and R. D. Goldman, ‘Vimentin induces changes in cell shape, motility, and adhesion during the epithelial to mesenchymal transition’, FASEB J. Off. Publ. Fed. Am. Soc. Exp. Biol., vol. 24, no. 6, pp. 1838–1851, June 2010, doi: 10.1096/fj.09-151639.

[7] Y. Jiu et al., ‘Bidirectional Interplay between Vimentin Intermediate Filaments and Contractile Actin Stress Fibers’, Cell Rep., vol. 11, no. 10, pp. 1511–1518, June 2015, doi: 10.1016/j.celrep.2015.05.008.

[8] K. Pogoda and P. A. Janmey, ‘Transmit and protect: the mechanical functions of intermediate filaments’, Curr. Opin. Cell Biol., vol. 85, p. 102281, Dec. 2023, doi: 10.1016/j.ceb.2023.102281.

[9] A.-B. Ndiaye, G. H. Koenderink, and M. Shemesh, ‘Intermediate Filaments in Cellular Mechanoresponsiveness: Mediating Cytoskeletal Crosstalk From Membrane to Nucleus and Back’, Front. Cell Dev. Biol., vol. 10, p. 882037, 2022, doi: 10.3389/fcell.2022.882037.

[10] J. Hu et al., ‘High stretchability, strength, and toughness of living cells enabled by hyperelastic vimentin intermediate filaments’, Proc. Natl. Acad. Sci., vol. 116, no. 35, pp. 17175–17180, Aug. 2019, doi: 10.1073/pnas.1903890116.

[11] L. Kreplak, H. Bär, J. F. Leterrier, H. Herrmann, and U. Aebi, ‘Exploring the mechanical behavior of single intermediate filaments’, J. Mol. Biol., vol. 354, no. 3, pp. 569–577, Dec. 2005, doi: 10.1016/j.jmb.2005.09.092.

[12] P. A. Janmey, U. Euteneuer, P. Traub, and M. Schliwa, ‘Viscoelastic properties of vimentin compared with other filamentous biopolymer networks’, J. Cell Biol., vol. 113, no. 1, pp. 155–160, Apr. 1991, doi: 10.1083/jcb.113.1.155.

13. ‘Vimentin intermediate filaments as structural and mechanical coordinators of mesenchymal cells | Nature Cell Biology’. Accessed: Jan. 06, 2026. [Online]. Available: https://www.nature.com/articles/s41556-025-01713-x

[14] Z. Gan et al., ‘Vimentin Intermediate Filaments Template Microtubule Networks to Enhance Persistence in Cell Polarity and Directed Migration’, Cell Syst., vol. 3, no. 3, pp. 252–263.e8, Sept. 2016, doi: 10.1016/j.cels.2016.08.007.

[15] C. J. Redmond and P. A. Coulombe, ‘Intermediate filaments as effectors of differentiation’, Curr. Opin. Cell Biol., vol. 68, pp. 155–162, Feb. 2021, doi: 10.1016/j.ceb.2020.10.009.

[16] J. Forsting, J. Kraxner, H. Witt, A. Janshoff, and S. Köster, ‘Vimentin Intermediate Filaments Undergo Irreversible Conformational Changes during Cyclic Loading’, Nano Lett., vol. 19, no. 10, pp. 7349–7356, Oct. 2019, doi: 10.1021/acs.nanolett.9b02972.

[17] C. G. Lopez, O. Saldanha, K. Huber, and S. Köster, ‘Lateral association and elongation of vimentin intermediate filament proteins: A time-resolved light-scattering study’, Proc. Natl. Acad. Sci. U. S. A., vol. 113, no. 40, pp. 11152–11157, Oct. 2016, doi: 10.1073/pnas.1606372113.

[18] M. Eibauer et al., ‘Vimentin filaments integrate low-complexity domains in a complex helical structure’, Nat. Struct. Mol. Biol., vol. 31, no. 6, pp. 939–949, June 2024, doi: 10.1038/s41594-024-01261-2.

[19] S. Winheim et al., ‘Deconstructing the Late Phase of Vimentin Assembly by Total Internal Reflection Fluorescence Microscopy (TIRFM)’, PLOS ONE, vol. 6, no. 4, p. e19202, Apr. 2011, doi: 10.1371/journal.pone.0019202.

[20] U. Rölleke, P. Kumari, R. Meyer, and S. Köster, ‘The unique biomechanics of intermediate filaments – From single filaments to cells and tissues’, Curr. Opin. Cell Biol., vol. 85, p. 102263, Dec. 2023, doi: 10.1016/j.ceb.2023.102263.

[21] R. Windoffer et al., ‘Quantitative mapping of keratin networks in 3D’, eLife, vol. 11, p. e75894, Feb. 2022, doi: 10.7554/eLife.75894.

[22] H. Iwatsuki and M. Suda, ‘Seven kinds of intermediate filament networks in the cytoplasm of polarized cells: structure and function’, Acta Histochem. Cytochem., vol. 43, no. 2, pp. 19–31, May 2010, doi: 10.1267/ahc.10009.

[23] H. Bloemendal and F. R. Pieper, ‘Intermediate filaments: known structure, unknown function’, Biochim. Biophys. Acta, vol. 1007, no. 3, pp. 245–253, Apr. 1989, doi: 10.1016/0167-4781(89)90144-9.

[24] D. A. Fletcher and R. D. Mullins, ‘Cell mechanics and the cytoskeleton’, Nature, vol. 463, no. 7280, pp. 485–492, Jan. 2010, doi: 10.1038/nature08908.

[25] A.-C. Reymann, J.-L. Martiel, T. Cambier, L. Blanchoin, R. Boujemaa-Paterski, and M. Théry, ‘Nucleation geometry governs ordered actin networks structures’, Nat. Mater., vol. 9, no. 10, pp. 827–832, Oct. 2010, doi: 10.1038/nmat2855.

[26] A.-C. Reymann et al., ‘Actin network architecture can determine myosin motor activity’, Science, vol. 336, no. 6086, pp. 1310–1314, June 2012, doi: 10.1126/science.1221708.

[27] R. Subramanian and T. M. Kapoor, ‘Building Complexity: Insights into Self-organized Assembly of Microtubule-based Architectures’, Dev. Cell, vol. 23, no. 5, pp. 874–885, Nov. 2012, doi: 10.1016/j.devcel.2012.10.011.

[28] F. Huber, A. Boire, M. P. López, and G. H. Koenderink, ‘Cytoskeletal crosstalk: when three different personalities team up’, Curr. Opin. Cell Biol., vol. 32, pp. 39–47, Feb. 2015, doi: 10.1016/j.ceb.2014.10.005.

[29] L. Pradeau-Phélut and S. Etienne-Manneville, ‘Cytoskeletal crosstalk: A focus on intermediate filaments’, Curr. Opin. Cell Biol., vol. 87, p. 102325, Apr. 2024, doi: 10.1016/j.ceb.2024.102325.

[30] L. Schaedel, C. Lorenz, A. V. Schepers, S. Klumpp, and S. Köster, ‘Vimentin intermediate filaments stabilize dynamic microtubules by direct interactions’, Nat. Commun., vol. 12, no. 1, p. 3799, June 2021, doi: 10.1038/s41467-021-23523-z.

[31] A. Aufderhorst-Roberts and G. H. Koenderink, ‘Stiffening and inelastic fluidization in vimentin intermediate filament networks’, Soft Matter, vol. 15, no. 36, pp. 7127–7136, Sept. 2019, doi: 10.1039/c9sm00590k.

32. ‘From mechanical resilience to active material properties in biopolymer networks | Nature Reviews Physics’. Accessed: Jan. 06, 2026. [Online]. Available: https://www.nature.com/articles/s42254-019-0036-4

[33] Y.-C. Lin, N. Y. Yao, C. P. Broedersz, H. Herrmann, F. C. Mackintosh, and D. A. Weitz, ‘Origins of elasticity in intermediate filament networks’, Phys. Rev. Lett., vol. 104, no. 5, p. 058101, Feb. 2010, doi: 10.1103/PhysRevLett.104.058101.

[34] A. V. Schepers, C. Lorenz, P. Nietmann, A. Janshoff, S. Klumpp, and S. Köster, ‘Multiscale mechanics and temporal evolution of vimentin intermediate filament networks’, Proc. Natl. Acad. Sci. U. S. A., vol. 118, no. 27, p. e2102026118, July 2021, doi: 10.1073/pnas.2102026118.

[35] S. Zafari, et al., ‘Vimentin networks at high strains’, Oct. 06, 2025, bioRxiv. doi: 10.1101/2025.10.06.680657.

[36] K. Bhattacharyya and S. Klumpp, ‘The unexpected structure and dynamics of vimentin networks’, J. Cell Biol., vol. 224, no. 4, Apr. 2025, doi: 10.1083/jcb.202503012.

[37] S. Chakraborty, J. Mahamid, and W. Baumeister, ‘Cryoelectron Tomography Reveals Nanoscale Organization of the Cytoskeleton and Its Relation to Microtubule Curvature Inside Cells’, Struct. Lond. Engl. 1993, vol. 28, no. 9, pp. 991–1003.e4, Sept. 2020, doi: 10.1016/j.str.2020.05.013.

[38] R. A, H. H, D. Mw, and G. Vi, ‘Microtubule-dependent transport of vimentin filament precursors is regulated by actin and by the concerted action of Rho- and p21-activated kinases’, FASEB J. Off. Publ. Fed. Am. Soc. Exp. Biol., vol. 28, no. 7, July 2014, doi: 10.1096/fj.14-250019.

[39] A. S. Jijumon et al., ‘Lysate-based pipeline to characterize microtubule-associated proteins uncovers unique microtubule behaviours’, Nat. Cell Biol., vol. 24, no. 2, pp. 253–267, Feb. 2022, doi: 10.1038/s41556-021-00825-4.

[40] A. S. Jijumon, A. Krishnan, and C. Janke, ‘A Platform for Medium-Throughput Cell-Free Analyses of Microtubule-Interacting Proteins Using Mammalian Cell Lysates’, Curr. Protoc., vol. 4, no. 6, p. e1070, June 2024, doi: 10.1002/cpz1.1070.

[41] M. Inada, G. Izawa, W. Kobayashi, and M. Ozawa, ‘293 cells express both epithelial as well as mesenchymal cell adhesion molecules’, Int. J. Mol. Med., vol. 37, no. 6, pp. 1521–1527, June 2016, doi: 10.3892/ijmm.2016.2568.

[42] H. Herrmann, L. Kreplak, and U. Aebi, ‘Isolation, characterization, and in vitro assembly of intermediate filaments’, Methods Cell Biol., vol. 78, pp. 3–24, 2004, doi: 10.1016/s0091-679x(04)78001-2.

[43] B. B. Mandelbrot, The fractal geometry of nature. San Francisco : W.H. Freeman, 1983. Accessed: Jan. 06, 2026. [Online]. Available: http://archive.org/details/fractalgeometryo00beno

[44] T. T. Falzone, M. Lenz, D. R. Kovar, and M. L. Gardel, ‘Assembly kinetics determine the architecture of α-actinin crosslinked F-actin networks’, Nat. Commun., vol. 3, p. 861, May 2012, doi: 10.1038/ncomms1862.

[45] C. Mencarelli, S. Ciolfi, D. Caroti, P. Lupetti, and R. Dallai, ‘Isomin: a novel cytoplasmic intermediate filament protein from an arthropod species’, BMC Biol., vol. 9, p. 17, Feb. 2011, doi: 10.1186/1741-7007-9-17.

[46] H. Herrmann and S. V. Strelkov, ‘History and phylogeny of intermediate filaments: now in insects’, BMC Biol., vol. 9, p. 16, Feb. 2011, doi: 10.1186/1741-7007-9-16.

[47] H. Drechsler, Y. Xu, V. F. Geyer, Y. Zhang, and S. Diez, ‘Multivalent electrostatic microtubule interactions of synthetic peptides are sufficient to mimic advanced MAP-like behavior’, Mol. Biol. Cell, vol. 30, no. 24, pp. 2953–2968, Nov. 2019, doi: 10.1091/mbc.E19-05-0247.

[48] H. Wu, Y. Shen, D. Wang, H. Herrmann, R. D. Goldman, and D. A. Weitz, ‘Effect of Divalent Cations on the Structure and Mechanics of Vimentin Intermediate Filaments’, Biophys. J., vol. 119, no. 1, pp. 55–64, July 2020, doi: 10.1016/j.bpj.2020.05.016.

[49] Q. D. Tran, V. Sorichetti, G. Pehau-Arnaudet, M. Lenz, and C. Leduc, ‘Fragmentation and Entanglement Limit Vimentin Intermediate Filament Assembly’, Phys. Rev. X, vol. 13, no. 1, p. 011014, Feb. 2023, doi: 10.1103/PhysRevX.13.011014.

[50] A. E. Patteson et al., ‘Vimentin protects cells against nuclear rupture and DNA damage during migration’, J. Cell Biol., vol. 218, no. 12, pp. 4079–4092, Dec. 2019, doi: 10.1083/jcb.201902046.

[51] F. K. Gyoeva and V. I. Gelfand, ‘Coalignment of vimentin intermediate filaments with microtubules depends on kinesin’, Nature, vol. 353, no. 6343, pp. 445–448, Oct. 1991, doi: 10.1038/353445a0.

[52] G. Gurland and G. G. Gundersen, ‘Stable, detyrosinated microtubules function to localize vimentin intermediate filaments in fibroblasts’, J. Cell Biol., vol. 131, no. 5, pp. 1275–1290, Dec. 1995, doi: 10.1083/jcb.131.5.1275.

[53] S.-J. Yao et al., ‘Surface keratin 1, a tumor-selective peptide target in human triple-negative breast cancer’, Sci. Rep., vol. 15, no. 1, p. 21644, July 2025, doi: 10.1038/s41598-025-05351-z.

[54] J. E. Eriksson et al., ‘Specific in vivo phosphorylation sites determine the assembly dynamics of vimentin intermediate filaments’, J. Cell Sci., vol. 117, no. Pt 6, pp. 919–932, Feb. 2004, doi: 10.1242/jcs.00906.

[55] N. T. Snider and M. B. Omary, ‘Post-translational modifications of intermediate filament proteins: mechanisms and functions’, Nat. Rev. Mol. Cell Biol., vol. 15, no. 3, pp. 163–177, Mar. 2014, doi: 10.1038/nrm3753.

[56] F. Danielsson, M. K. Peterson, H. Caldeira Araújo, F. Lautenschläger, and A. K. B. Gad, ‘Vimentin Diversity in Health and Disease’, Cells, vol. 7, no. 10, p. 147, Sept. 2018, doi: 10.3390/cells7100147.

[57] J. Ivaska, H.-M. Pallari, J. Nevo, and J. E. Eriksson, ‘Novel functions of vimentin in cell adhesion, migration, and signaling’, Exp. Cell Res., vol. 313, no. 10, pp. 2050–2062, June 2007, doi: 10.1016/j.yexcr.2007.03.040.

[58] A. A. Challa and B. Stefanovic, ‘A novel role of vimentin filaments: binding and stabilization of collagen mRNAs’, Mol. Cell. Biol., vol. 31, no. 18, pp. 3773–3789, Sept. 2011, doi: 10.1128/MCB.05263-11.

[59] K. M. Ridge, J. E. Eriksson, M. Pekny, and R. D. Goldman, ‘Roles of vimentin in health and disease’, Genes Dev., vol. 36, no. 7–8, pp. 391–407, Apr. 2022, doi: 10.1101/gad.349358.122.

[60] B. Eckes et al., ‘Impaired mechanical stability, migration and contractile capacity in vimentin-deficient fibroblasts’, J. Cell Sci., vol. 111 ( Pt 13), pp. 1897–1907, July 1998, doi: 10.1242/jcs.111.13.1897.

[61] J. Block et al., ‘Viscoelastic properties of vimentin originate from nonequilibrium conformational changes’, Sci. Adv., vol. 4, no. 6, p. eaat1161, June 2018, doi: 10.1126/sciadv.aat1161.

[62] B. Nöding and S. Köster, ‘Intermediate filaments in small configuration spaces’, Phys. Rev. Lett., vol. 108, no. 8, p. 088101, Feb. 2012, doi: 10.1103/PhysRevLett.108.088101.

[63] J. Block et al., ‘Nonlinear Loading-Rate-Dependent Force Response of Individual Vimentin Intermediate Filaments to Applied Strain’, Phys. Rev. Lett., vol. 118, no. 4, p. 048101, Jan. 2017, doi: 10.1103/PhysRevLett.118.048101.

[64] N. Mücke et al., ‘Molecular and biophysical characterization of assembly-starter units of human vimentin’, J. Mol. Biol., vol. 340, no. 1, pp. 97–114, June 2004, doi: 10.1016/j.jmb.2004.04.039.

[65] B. Morgan et al., ‘Real-time monitoring of basal H2O2 levels with peroxiredoxin-based probes’, Nat. Chem. Biol., vol. 12, no. 6, pp. 437–443, June 2016, doi: 10.1038/nchembio.2067.

[66] J. Schindelin et al., ‘Fiji: an open-source platform for biological-image analysis’, Nat. Methods, vol. 9, no. 7, pp. 676–682, July 2012, doi: 10.1038/nmeth.2019.

[67] J. Hoshen and R. Kopelman, ‘Percolation and cluster distribution. I. Cluster multiple labeling technique and critical concentration algorithm’, Phys. Rev. B, vol. 14, no. 8, pp. 3438–3445, Oct. 1976, doi: 10.1103/PhysRevB.14.3438.

68. [68] Theory of Simple Liquids. 2006. Accessed: Jan. 06, 2026. [Online]. Available: https://shop.elsevier.com/books/theory-of-simple-liquids/hansen/978-0-12-370535-8

[69] M. R. Shaebani, H. Rieger, and Z. Sadjadi, ‘Kinematics of persistent random walkers with two distinct modes of motion’, *Phys*. Rev. E, vol. 106, no. 3, p. 034105, Sept. 2022, doi: 10.1103/PhysRevE.106.034105.

