## Supplementary_figures for "Cytosolic factors govern vimentin network architecture and mechanics"

### Supplementary Figure 1

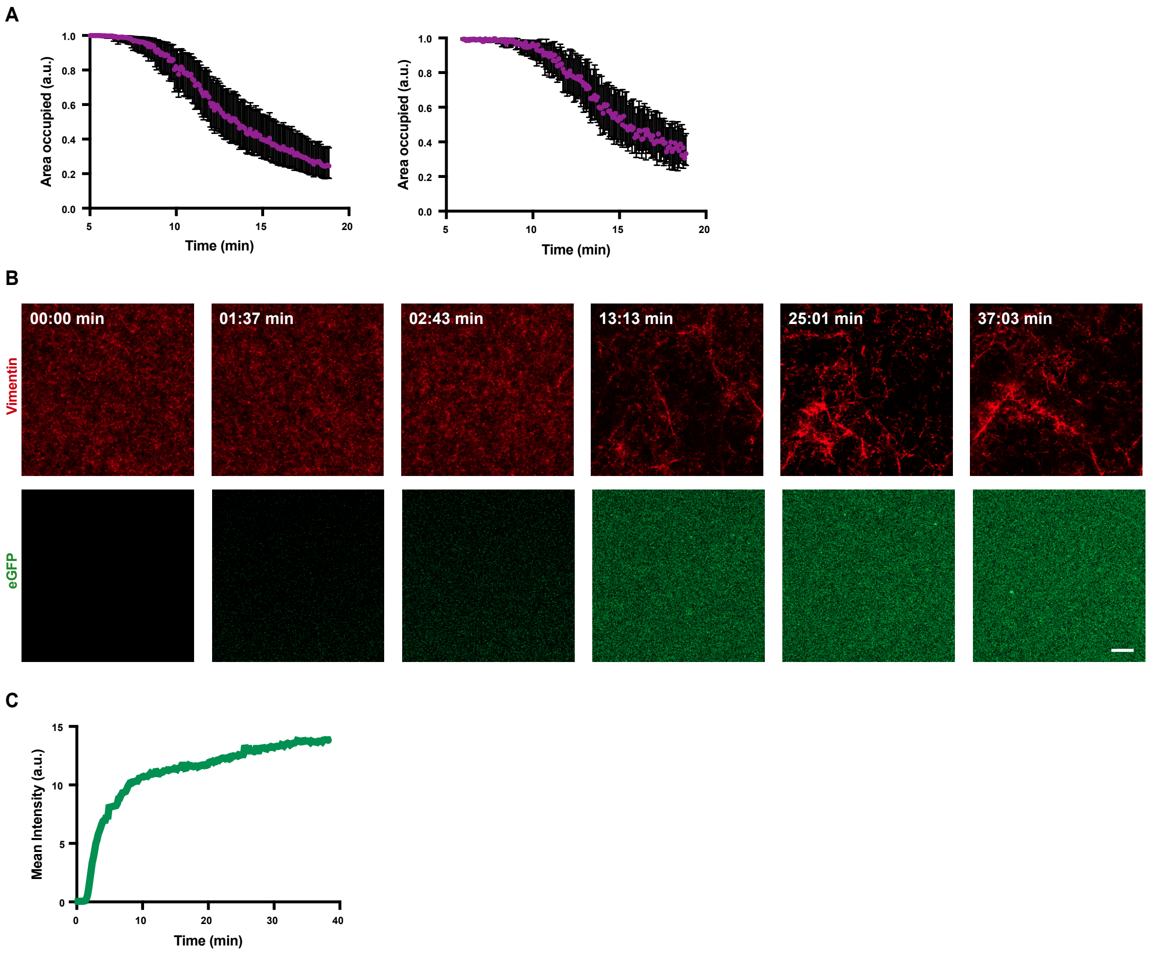


**Supplementary Figure 1: (A)** Plots showing time evolution of occupied area fraction from additional replicates. Time-lapse imaging was done as mentioned in the experimental setup shown in Fig.1J and occupied area was derived using the analysis pipeline (binary conversion) shown in Fig.1I. The error bar indicates the variation range of the area fraction for the admissible range of intensity thresholds over time. **(B)** Representative images showing time evolution of the vimentin network formation in the presence of eGFP. Here eGFP acts as a dummy protein in the cell extract. Network formation roughly relates to the diffusion of the cell extract into the FoV **(C)** Plot showing time evolution of mean intensity of eGFP. Mean intensity of the FoV was measured in Fiji [66]. Scale bar = 10 µm

### Supplementary Figure 2

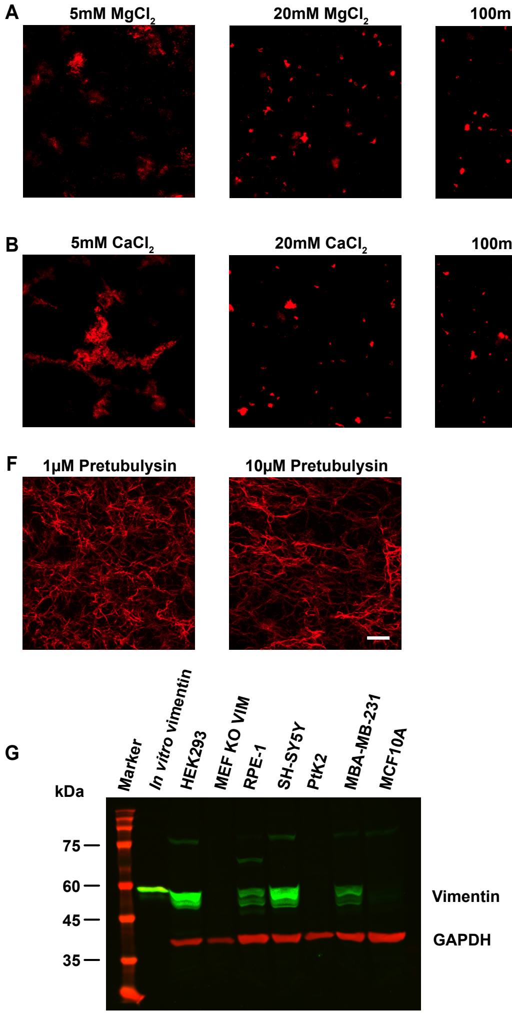


**Supplementary Figure 2:** Evidence supporting involvement of a cellular factor. **(A)** Representative images showing vimentin in the presence of varying concentrations (5mM, 20mM, 100mM) of $MgCl_2$. **(B)** Representative images showing vimentin in the presence of varying concentrations (5mM, 20mM, 100mM) of $CaCl_2$. **(C)** Representative image showing vimentin in the presence of denatured cell extract. **(D)** Representative image showing vimentin network in crude extract. **(E)** Representative image showing vimentin tetramers (0.2mg/ml) in the presence of cell extract (4mg/ml). **(F)** Representative images showing vimentin network in cell extract (4mg/ml) containing 1µM and 10µM of Pretubulysin (microtubule depolymerizing drug). **(G)** Representative immunoblot showing endogenous vimentin expression in HEK293, MEF VIM KO, RPE-1, SH-SY5Y, PtK2, MDA-MB-231 and MCF10A cell extracts. In all these experiments, vimentin filaments were assembled at 0.2mg/mL and further diluted to 0.04mg/mL with the respective cell extract. Experiments were performed in three independent biological replicates. Scale bar = 10 µm

### Supplementary Figure 3

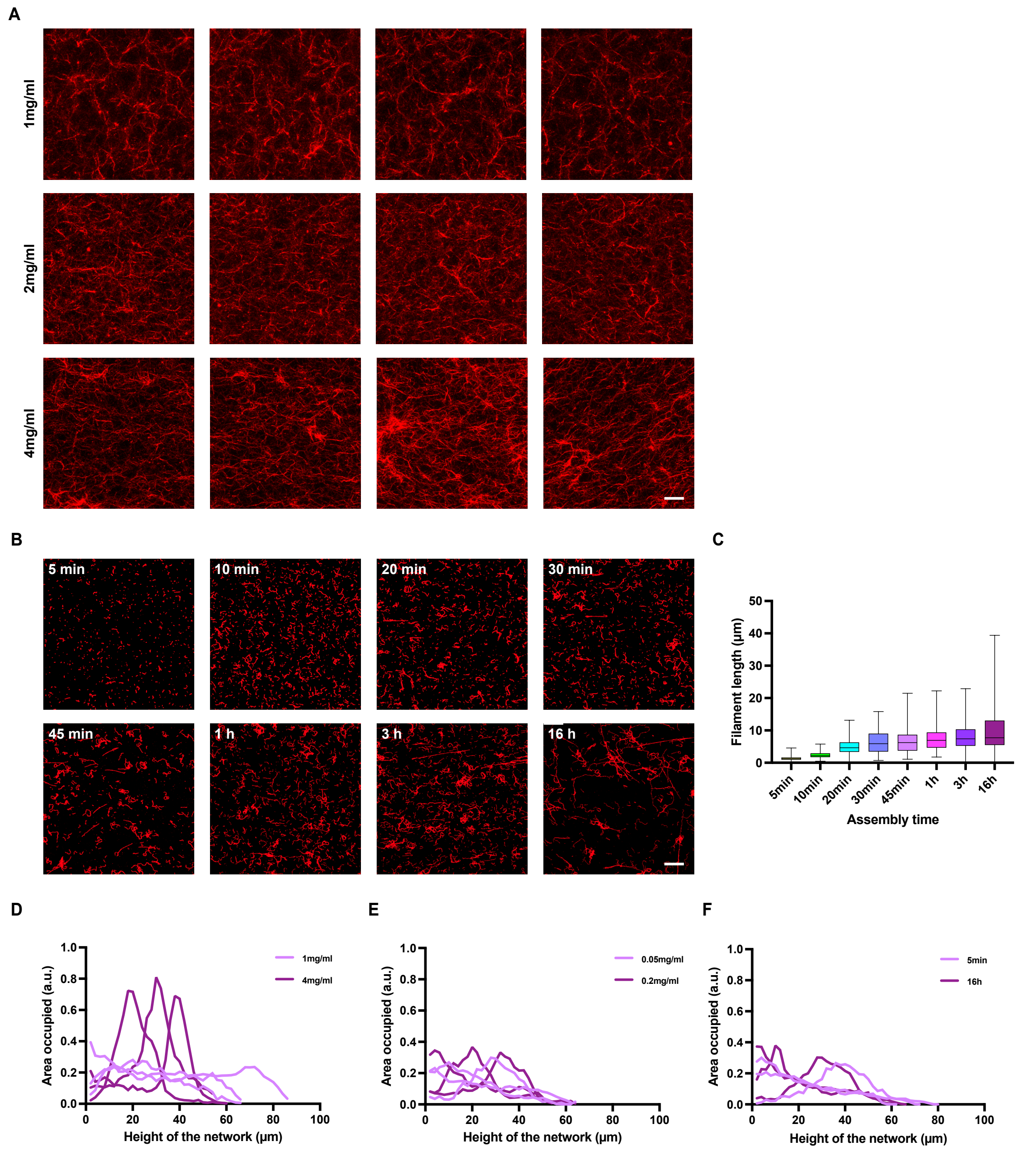


**Supplementary Figure 3**: **(A)** Additional examples of images showing vimentin network with varying concentration of the cell extract (1mg/ml, 2mg/ml, 4mg/ml). **(B)** Temporal evolution of vimentin filament length. Representative images at different time points after start of assembly. Vimentin filaments were assembled at 0.2mg/ml. Samples were imaged 5 mins after initiation of the assembly and the experiment ran till 16h. **(C)** Plot showing traced filament lengths at different time points after starting the assembly. Filaments were manually traced in Fiji [66]. Boxplots include the median as the center line, the 25th and 75th percentiles as box limits and the entire data range as whiskers. For each time point, 200 filaments were measured which were pooled from two replicates. **(D, E, F)** Plots showing the area occupied fraction in the vertical range of the vimentin network spanning the flow chamber; In E and F extract concentration was kept at 1mg/ml. Scale bar = 10 µm.

### Supplementary Figure 4

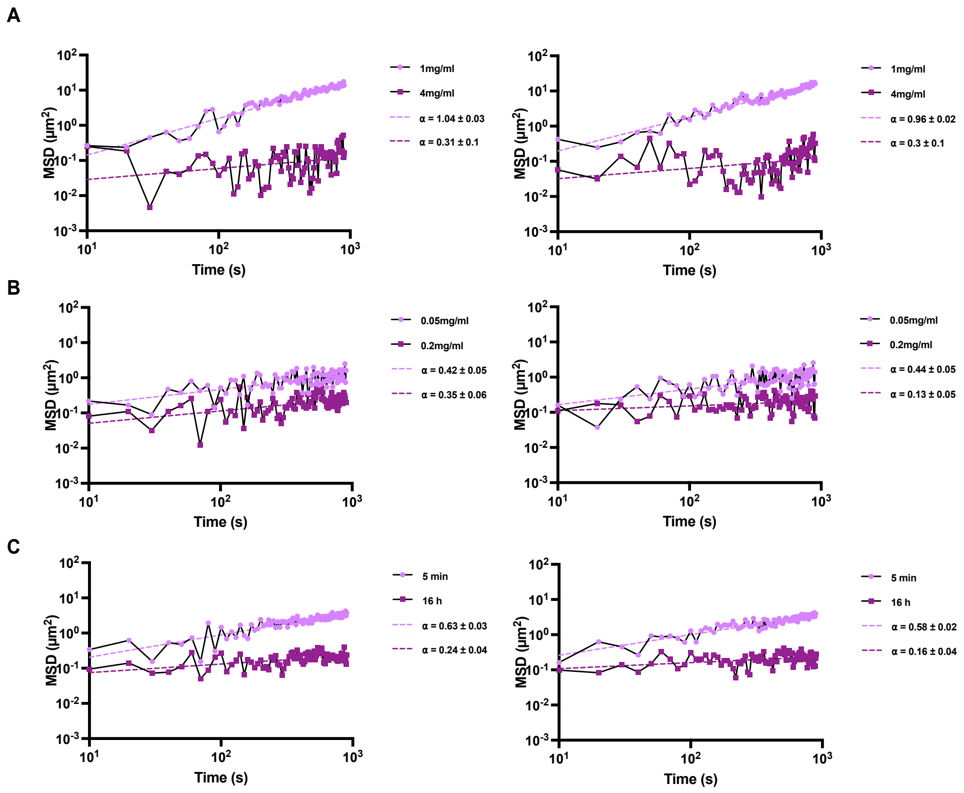


**Supplementary Figure 4**: Additional replicates for the analysis of MSD. **(A)** Plot showing the mean-squared displacement (MSD) of the centre of mass of vimentin networks with varying extract concentrations. **(B)** Plot showing the MSD of the centre of mass of vimentin networks with varying *in vitro* vimentin concentrations. **(C)** Plot showing the MSD of the centre of mass of the vimentin network fluctuations with varying assembly time of the *in vitro* assembled vimentin. The centre of mass rearrangements follow an anomalous diffusion $MSD(\Delta t) \propto \Delta t^{\alpha}$. Here, α is the anomalous exponent, which reflects the overall dynamics of the network: sub-diffusion (slower than diffusion) for $0<\alpha<1$, normal diffusion for $\alpha=1$, and super-diffusion (faster than diffusion) for $1<\alpha<2$; For a detailed explanation on the quantification of relative network fluctuations and MSD, refer to the methods section.

**Supplementary Movie 1:** Time evolution of the vimentin network formation. Time-lapse imaging was done as mentioned in the experimental setup shown in Fig.1J. Scale bar = 10 µm
